# DDI-GPT: Explainable Prediction of Drug-Drug Interactions using Large Language Models enhanced with Knowledge Graphs

**DOI:** 10.1101/2024.12.06.627266

**Authors:** Chengqi Xu, Krishna C. Bulusu, Heng Pan, Olivier Elemento

## Abstract

Identifying potential drug-drug interactions (DDIs) before clinical use is essential for patient safety yet remains a significant challenge in drug development. We presented DDI-GPT, a deep learning framework that predicts DDIs by combining knowledge graphs (KGs) and pre-trained large language models (LLMs), enabling early detection of potential drug interactions. We demonstrated that DDI-GPT outperforms current state-of-the-art methods by capturing contextual dependencies between biomedical entities to infer potential DDIs. Through feature attribution methods, we show that our explainable deep learning (DL) models enhance the quality of explanations on the pathways and interactome networks. Using TwoSIDES as our benchmark dataset, DDI-GPT achieved the best performance of 0.964 in AUROC compared with existing DL methods. We also applied DDI-GPT to perform zero-shot prediction on 9,480 DDI records, encompassing 442 distinct drugs from the FDA Adverse Event Reporting System. DDI-GPT can attain a high accuracy of in 0.84 AUROC, with an improvement of 14% compared to the best previously published method. We explored model interpretations on predicted DDIs involving Bruton’s tyrosine kinase (BTK) inhibitors and uncovered CYP3A-enriched signals underlying the contaminant use of BTK inhibitors with other drugs leading to toxicity. Altogether, DDI-GPT, implemented as both a web server platform and a software package, identifies DDI events and offers a deep learning tool for drug safety use with expandable features.

## Introduction

Drug-drug interactions (DDIs) critically impact patient safety in healthcare, accounting for approximately 1% of hospitalizations in the general population and 2-5% among elderly patients^1–3^. As combination therapies become increasingly prevalent for treating complex diseases like cancer, the risks of DDIs continue to rise^1^. The challenges of predicting DDIs are particularly evident in the treatment of diffuse large B-cell lymphoma (DLBCL). This most common form of non-Hodgkin lymphoma requires sophisticated treatment approaches due to its heterogeneous genetic and clinical profile^2^. Recent therapeutic advances have led to the incorporation of Bruton’s tyrosine kinase (BTK) inhibitors alongside standard chemotherapy regimens, showing promising results for DLBCL patients^3^. However, this combination therapy introduces new risks of adverse drug interactions. Clinical studies have revealed serious complications, including heightened bleeding risk when BTK inhibitors are administered with anticoagulants or antiplatelet agents ^4,5^. Moreover, when patients receive BTK inhibitors alongside strong CYP3A4 inhibitors, the resulting elevated drug concentrations can lead to significant toxicity^6,7^. These interactions, typically discovered only after drugs enter clinical use through healthcare provider reports, underscore the critical need for predictive DDI screening before clinical implementation.

Recent technological advances enable DDI prediction through systematic analysis of biomedical knowledge graphs (KGs). Over the past decades, researchers have constructed large KGs such as Hetionet, DRKG, and iBKH ^8–11^ through literature mining and database integration, establishing comprehensive maps of relationships among drugs, proteins, diseases, and other biological entities. These resources provide a rich foundation for identifying DDIs by capturing both entity attributes and network topology. Previous work that combined KGs with traditional machine learning such as tensor factorization-based method faced common limitations because they cannot maintain the distinct types of relationships embedded within the KG^12^.

Deep learning models, particularly graph neural networks (GNNs), have emerged as the leading approach for analyzing heterogenous graph in DDI predictions^13,14^. However, GNNs face significant limitations when encoding textual information from KGs. They typically rely on context-independent shallow embeddings, where each entity is represented by a single vector regardless of its biological context. This approach creates two key challenges: biological entities must share the same representation even when their meaning varies across contexts, and the embeddings become highly sensitive to missing relationships in the graph structure ^15–17^. Moreover, existing models have not been validated on independent and up-to-date datasets, making it questionable of their abilities to deal with new or understudied drugs that lack DDI records.

To address these questions, we introduce DDI-GPT, a deep learning model that leverages the LLM framework to predict DDIs. Our curated KG incorporates comprehensive biomedical knowledge, including drug targets, biological pathways, anatomical therapeutic chemicals, and drug categories, creating a rich feature space for DDI prediction. Unlike GNNs that rely primarily on shallow embeddings and graph topologies, DDI-GPT utilizes semantic comprehension from extensive biomedical corpora to enhance prediction accuracy. When the KG contains missing links, DDI-GPT can infer potential interactions between entities. DDI-GPT was trained and benchmarked on textual data between drug pairs from the TwoSIDES dataset^18^. We translate drug-related KG information into text, which we call a sentence tree, converting the nodes and edges of the KG into natural language descriptions that capture drug relationships. Once trained, DDI-GPT can predict new DDIs by incorporating entities from a new drug’s KG without requiring additional parameters or fine-tuning. We evaluated DDI-GPT on a newly curated independent real-world dataset from updated FDA labels. To understand the biological mechanisms underlying our predictions, we developed an evaluation framework that identifies the most influential genes driving each predicted interaction. As proof of concept, we applied this framework to analyze potential DDIs for acalabrutinib, a BTK inhibitor. Our evaluation module assigns importance scores to genes involved in each predicted interaction, providing transparent rationales for DDI-GPT’s predictions. We then integrated these scores with pathway enrichment analysis and calculated network-based distances between drug targets and genes associated with adverse reactions, enabling mechanistic insights into predicted DDIs.

## Results

### KG-informed deep learning for DDI predictions

We introduced DDI-GPT, a novel deep learning model that predicts DDI for pairs of drugs to potentially be used in combination. DDI-GPT analyzes drug pairs along with their associated biomedical information encoded in KGs (**Fig. 1a-f**) and outputs binary predictions based on interaction likelihood values. We classify these as "interacting" (>0.5) or "non-interacting" (<0.5) throughout this paper (**Methods**). The architecture comprises three primary modules: the knowledge module (**Fig. 1a-c**), the prediction module (**Fig. 1d**), and the evaluation module (**Fig. 1e-g**).

**Fig. 1:**
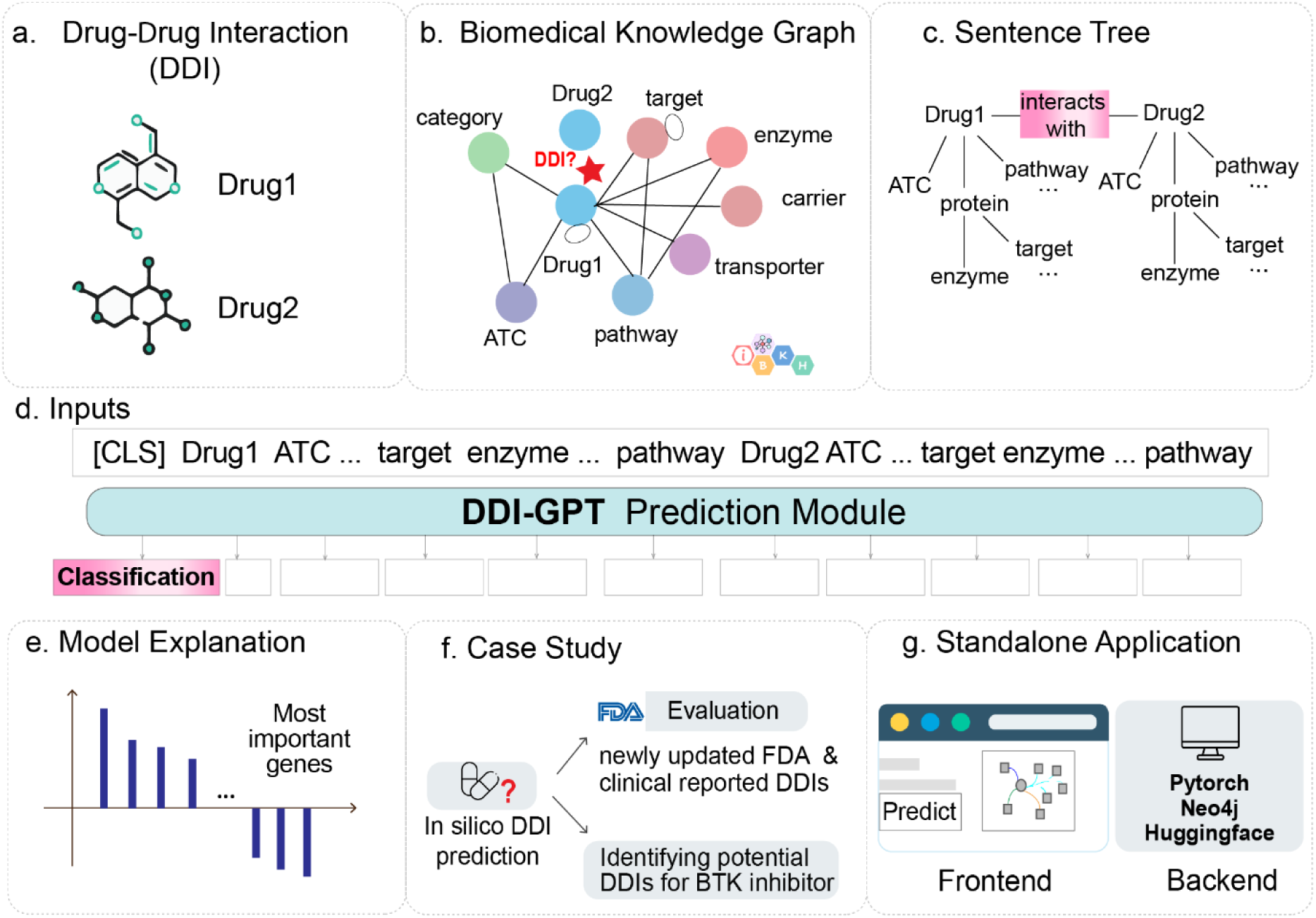
Overview of the DDI-GPT framework. **a.** Constructing drug-related multimodal representation through biomedical knowledge integration based on iBKH database. iBKH harmonizes data from 18 publicly available biomedical knowledge sources. We first collected diverse relation data which involves any drug entities in iBKH. Knowledge from diverse biomedical entities was integrated to build our curated drug-centered KG. **b.** We converted each DDI event into a sentence tree via knowledge injection from KG. **c.** The knowledge-enriched tree was translated into natural text and a task-specific prompt was created. The prompt was designed to generate binary class predictions of DDIs. **d.** A transformer-based prediction module was implemented to DDI-GPT for novel DDI predictions. **e.** We explained LLMs’ reasoning by ranking the most important genes nominated by DDI-GPT. **f.** We validated our model on a new reference dataset which includes newly updated FDA and clinically reported DDIs. As a proof of concept, we analyzed *in silico* DDI predictions for the BTK inhibitor acalabrutinib. **g.** An interactive, dynamic web server was developed, which allows users to perform on-demand interface predictions using the DDI-GPT framework.

The knowledge module processes drug pairs as input, identifies their neighborhood relationships in the KG, and transforms this information into a knowledge-rich sentence tree (**Fig. 2a**). To curate a KG specific for DDI prediction, we selected drug-associated information from the integrative Biomedical Knowledge Hub (iBKH), including information across multiple biological scales - from biological processes and molecular pathways to anatomical and phenotypic scales, as well as therapeutic mechanisms of action (**Fig. 2a**). iBKH is a curated KG designed to provide a comprehensive, multimodal view of biomedical entities^10^. Our final KG encompassed 129,361 entities, including targets, metabolized enzymes, carriers, transporters, small molecule pathways, ATC codes, and drug indication categories (**Supplementary Fig. 1a-b**), along with 4,033,682 relationships. DDI-GPT employs a novel approach using knowledge-enriched sentences to represent drug combinations. These sentences are converted into token-level embedding representations. We introduced a custom visibility matrix to mask attention between tokens belonging to different drugs. While standard self-attention mechanisms allow every token to attend to all other tokens regardless of their drug association, our approach preserves the original KG topology, maintaining valuable structural information about drug relationships. For example, Brevibloc (token 1) and beta-Adrenergic (token 5) are related, but should not interact with Calcium (token 21) (**Fig. 2b**).

**Fig. 2:**
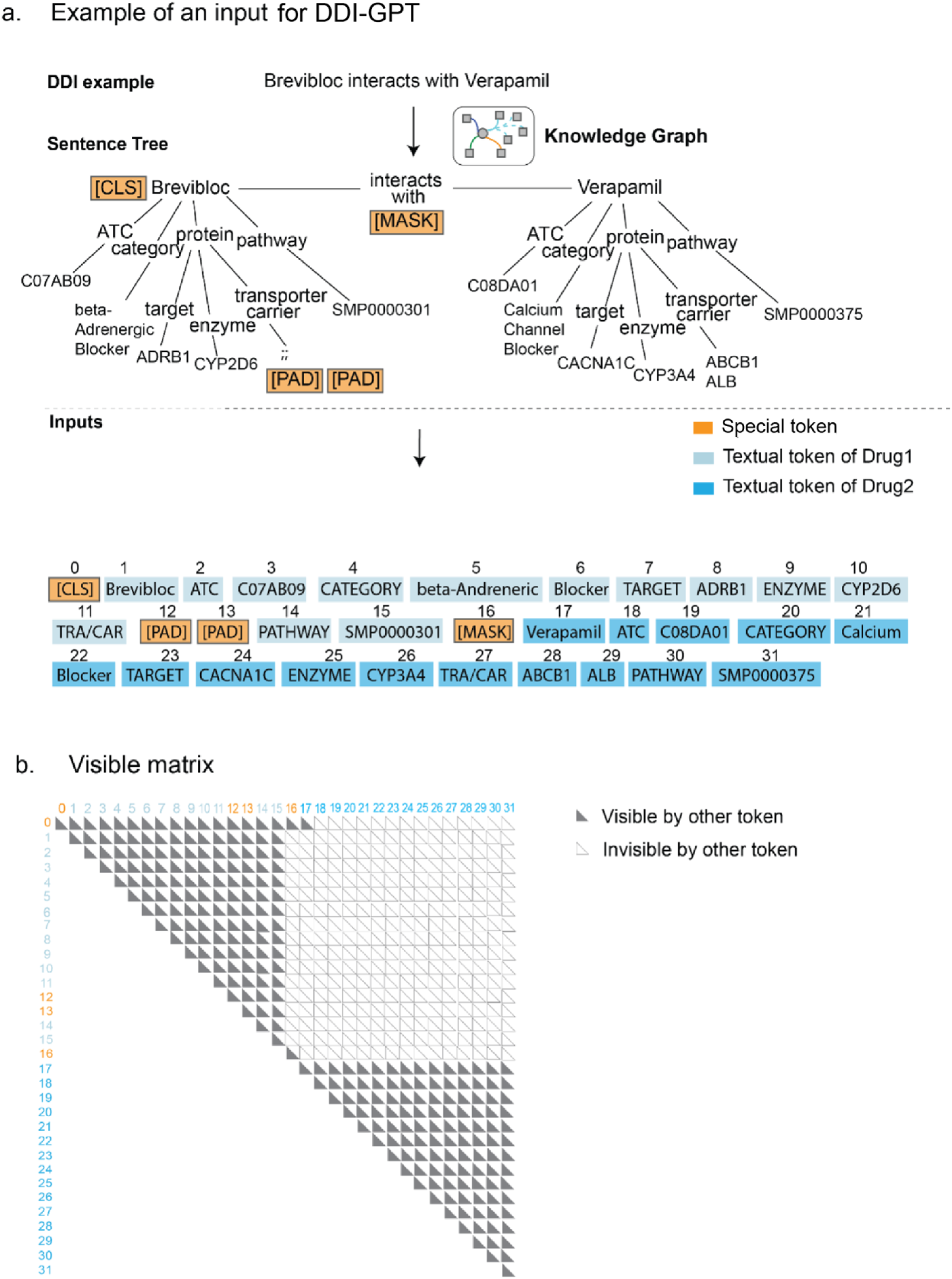
An example of knowledge graph is enhanced to construct inputs for DDI-GPT. **a.** The process of converting a DDI event, based on KG, into a sentence tree. Then the sentence tree is serialized into a natural text representation. This serialization process starts with a sentence containing the first drug’s ATC, category, target, enzyme, transporter/carrier (TRA/CAR), and pathway. It then describes the second drug. The special padded token PAD is added to empty position of a sentence to ensure the sentence length are same if there is absent feature. The special masked token MASK is an indicator to the model that DDI label should be neglected during the process of training. The special token CLS is used for classification of DDI label. **b.** The visible matrix limits the visible areas of each token by two rules: tokens belonging to two different drugs are masked; but tokens from the same drug are visible to each other.

The DDI-GPT prediction module consists of a GPT backbone optimized on 500,000 PubMed abstracts with an additional prediction layer for DDI classification^19^. We evaluated DDI-GPT using the expert-curated TwoSIDES dataset, which comprises 645 drugs with 63,437 interacting DDIs and 63,437 non-interacting DDIs randomly sampled from unseen pairs^18^. Through large-scale supervised training, DDI-GPT generates meaningful representations of all entities in the sentence tree, enabling transfer of DDI-associated knowledge from well-annotated drugs to those with limited interaction data.

DDI-GPT includes an evaluation module essential for understanding biological plausibility and building trust in model predictions. To help human experts understand the reasoning behind DDI predictions and validate model hypotheses, DDI-GPT’s explainer utilizes token importance scores to identify and represent critical drug-associated knowledge. For every token in the sentence tree, DDI-GPT generates an importance score between 0 and 1, relating tokens to DDI predictions through attention mechanisms. A score of 1 indicates the token is vital for prediction while 0 suggests it is irrelevant (**Fig. 2e**). We also applied DDI-GPT to test its zero-shot prediction capabilities, as validated on a new reference dataset including FDA and clinical DDI reports from 2013 to 2023, while the TwoSIDES training data predated 2013 (**Fig. 2f**). We developed an interactive tool that displays DDI-GPT’s predictions and interpretable scores associated with the input sentence tree (**Fig. 2g, Supplementary Fig. 3, code availability**), supporting real-world clinical decision-making.

### Benchmark evaluation of DDI-GPT

We implemented standard benchmarking to evaluate DDI prediction models using the TwoSIDES dataset. We randomly shuffled DDI pairs and set aside 20% (n=25,376) as a holdout set. We compared DDI-GPT against five state-of-the-art machine learning methods, including structural similarity approaches (*DeepDDI*), modeling with heterogeneous biomedical entities (*Decagon*), and advanced GNN methods such as weighted graph convolutional networks (*MR-GNN*), heterogeneous attention networks (*SSI-DDI*), and nonlinear encoder-decoder layers (*CASTER*)^15,20–23^.

We evaluated model performance using three metrics: the area under the receiver operating characteristic curve (AUROC), the area under the precision-recall curve (AUPRC), and Precision@50. All five state-of-the-art methods achieved AUROC values above 0.80, with CASTER performing best at 0.920 AUROC. DDI-GPT surpassed this performance with an AUROC of 0.964. For Precision@50, which assesses the accuracy of the top 50 ranked predictions for interacting DDIs, DDI-GPT achieved 0.890, representing a 16% improvement over existing benchmarks (**Table 1**).

**Table 1.** Validation of DDI-GPT and five state-of-the-art methods on the TwoSIDES dataset.

| Method | Representation | Architecture | AUROC | AUPRC | Precision @50 |
| --- | --- | --- | --- | --- | --- |
| DDI-GPT<br>(Our model) | Knowledge Graph<br>Augmented Sentence<br>Tree | GPT framework | **0.964±0.002** | **0.943±0.001** | **0.890±0.002** |
| DeepDDI<sup>19</sup> | Structural Similarity | Fully Connected<br>Neural Networks | 0.826±0.002 | 0.812±0.001 | 0.734±0.001 |
| Decagon<sup>14</sup> | Heterogenous<br>Knowledge Graph | Encoder and Decoder<br>Layers | 0.863±0.002 | 0.809±0.002 | 0.803±0.001 |
| MR-GNN<sup>20</sup> | Molecular Graph | Weighted Graph<br>Convolutional<br>Networks and LSTM | 0.845±0.003 | 0.8029±0.003 | 0.728±0.001 |
| SSI-DDI<sup>21</sup> | Chemical Substructure | Graph Attention<br>Network | 0.858±0.001 | 0.782±0.001 | 0.827±0.001 |
| CASTER<sup>22</sup> | Chemical Substructure | Encoder and Decoder<br>Layers | 0.920±0.005 | 0.935±0.001 | 0.887±0.002 |

To understand DDI-GPT’s predictive capabilities, we examined cutting-edge deep learning architectures commonly used in sentence classification. Convolutional Neural Networks (CNNs) employ convolutional filters to capture local features in text, while Recurrent Neural Networks (RNNs) process text sequentially, propagating information between words^12,16^. We applied these models to the TwoSIDES holdout dataset using identical inputs for model building and evaluation. Both CNN and RNN models showed inferior performance compared to DDI-GPT, highlighting how our unique deep learning architecture captures more information from the features than previous neural network approaches (**Table 2**).

**Table 2.** Prediction performance of DDI-GPT versus the alternative deep learning models using the four input drug features injected from a biomedical KG.

| Method | Drug features injected from biomedical Graph | Architecture | AUROC | AUPRC | Precision@50 |
| --- | --- | --- | --- | --- | --- |
| DDI-GPT (full KG) | ATC, category, pathway, targets | GPT framework | **0.964±0.002** | **0.943±0.001** | **0.890±0.002** |
| CNN (full KG) | ATC, category, pathway, targets | Convolutional Networks | 0.817±0.001 | 0.657±0.002 | 0.739±0.002 |
| RNN (full KG) | ATC, category, pathway, targets | Recurrent Neural Networks | 0.710±0.002 | 0.433±0.007 | 0.667±0.005 |

We next evaluated how varying the components of KG background knowledge affects drug representations and DDI predictions. We conducted ablation studies using three KGs of different sizes, based on the number of biomedical entities included during training: Full KG (complete enriched dataset), BP KG (biological processes only), and PP KG (pharmacological properties only) (**Supplementary Note 2**). We found that DDI-GPT with full KG features substantially outperformed both BP KG and PP KG variants (**Table 3**). Both DDI-GPT and other deep learning models (RNN, CNN) showed optimal performance when using the full KG as input (**Supplementary Table 2**), confirming that comprehensive KG representation of prior drug knowledge is critical for model performance. These improvements clearly demonstrate that our novel deep learning architecture and complete KG features significantly enhance DDI-GPT’s prediction accuracy.

**Table 3.**
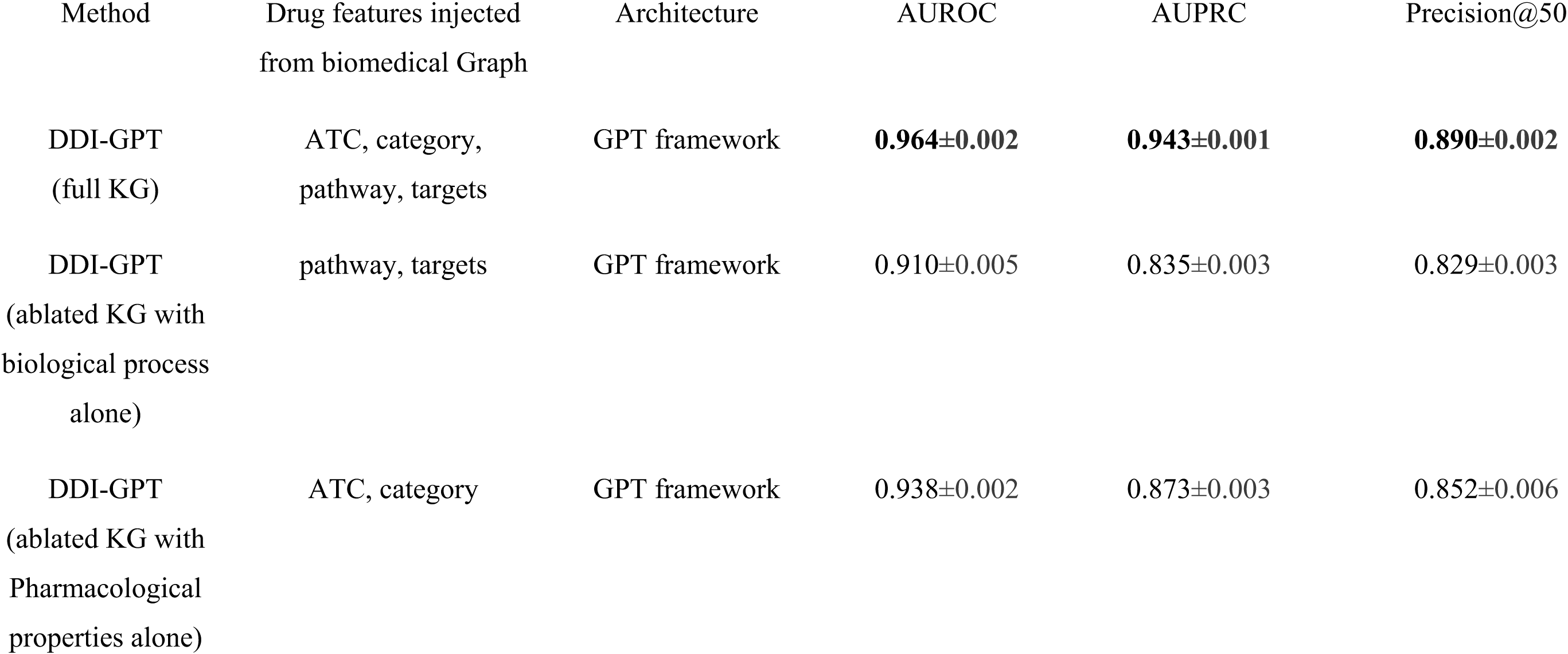
KG features are critical for model performance. On limited KG, DDI-GPT still achieves robust performance.

### Accurately predicting newly-updated DDIs from FDA data

Because drugs sharing similar mechanisms of action often exhibit overlapping DDI profiles, we evaluated DDI-GPT’s effectiveness on a validation dataset of newly updated DDIs. When models rely on superficial patterns rather than deeper mechanistic understanding, they often fail to extend predictions to drugs lacking experimental data or complete KG information. This ’shortcut learning’ phenomenon can lead to high performance on standard benchmarks but poor zero-shot prediction capability^24^. Therefore, a model’s ability to infer missing links in KGs and make reliable predictions for drugs without prior interaction data is fundamentally important

The TwoSIDES dataset used for training contained data collected prior to 2013. To assess our model’s ability to predict more recent DDIs, we collected newly reported DDI data from the FDA Adverse Event Reporting System (FAERS)^25^ and Micromedex Platform^26^ covering 2013Q1 to 2023Q2. From an initial set of 10,014 FAERS records of concomitant drug use, we performed rigorous filtering and validation to curate a high-confidence dataset (**Methods**). We benchmarked DDI-GPT against CASTER, the second-best performing model, on this new independent dataset. DDI-GPT significantly outperformed CASTER in zero-shot DDI predictions, showing improvements of 14% in AUROC and 15% in AUPRC.

### DDIs with acalabrutinib predicted by DDI-GPT are enriched in drugs metabolized by CYP3 family

After validating our model on an independent dataset, we applied DDI-GPT to predict potential DDIs for acalabrutinib, a recently FDA-approved drug. The next-generation Bruton’s tyrosine kinase (BTK) inhibitor acalabrutinib was approved after our TwoSIDES training dataset was released, making it an ideal candidate for evaluating zero-shot DDI prediction capabilities.

We performed a pairwise combinatorial assessment of 442 unique drugs from our independent dataset. Since acalabrutinib was absent from the initial training data, we updated the KG to incorporate comprehensive features for prediction purposes. Of the 442 drug combinations tested with acalabrutinib, 81 were predicted to have DDIs (aca-DDIs). We found that combinations involving CYP3A enzymes were enriched among DDI drugs with acalabrutinib (**Fig. 3a, Supplementary Fig. 2**). We further validated this finding through disproportionality analysis using the enzyme odds ratio (EOR) method, based on a 2×2 contingency table (**Supplementary Table 2, Methods**). The analysis demonstrated that drugs metabolized predominantly by CYP3A enzymes are 3.01 times more likely to cause DDI events when co-administered with acalabrutinib compared to drugs not metabolized by CYP3A (OR = 3.01 [1.57-5.78], P = 0.00043) (**Fig. 3b**).

**Fig. 3:**
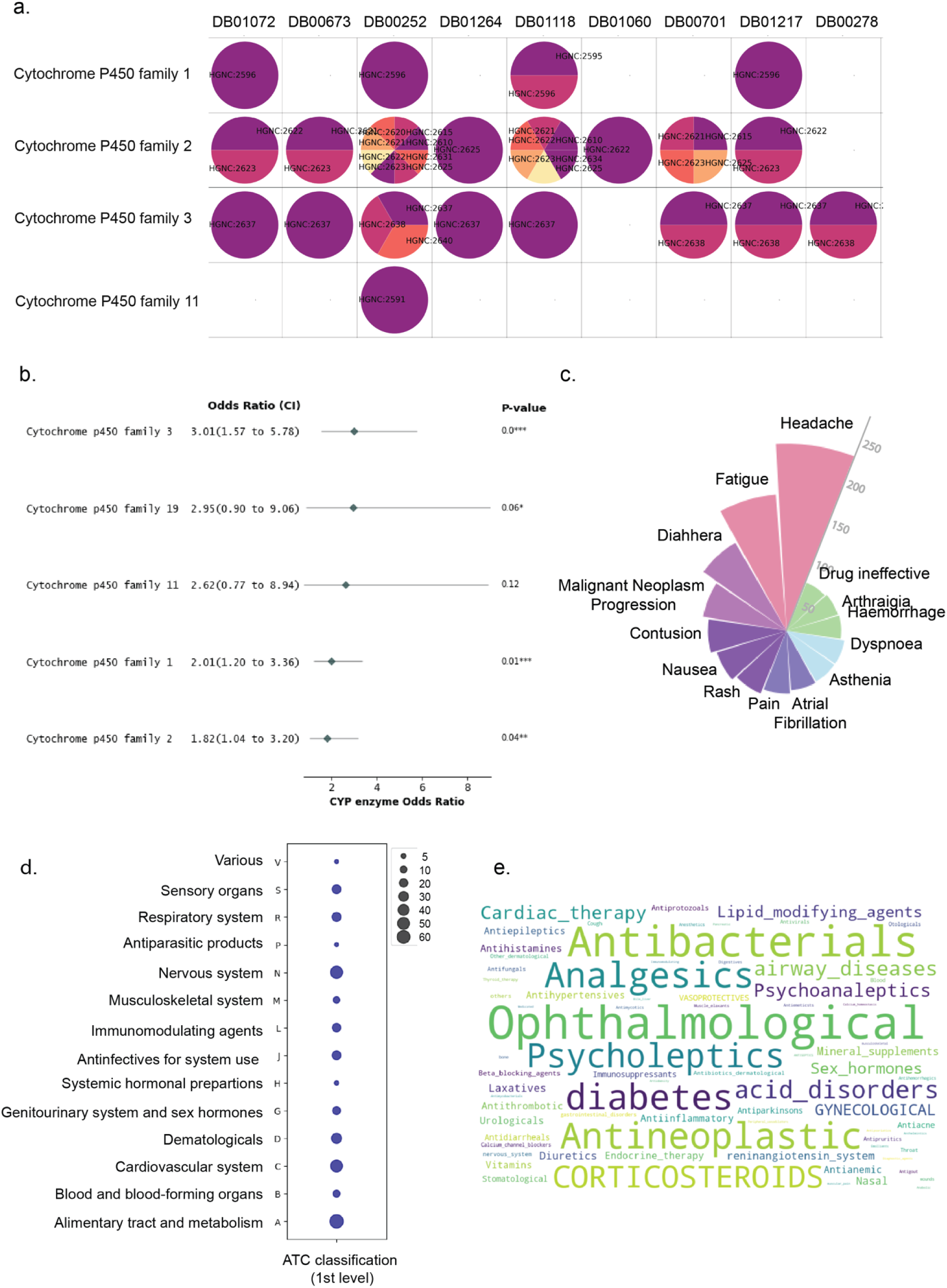
DDI-GPT-predicted aca-DDIs are associated with CYP3 enzyme family. **a.** Enzymes family of acalabrutinib predicted DDI drug for disproportional analysis in **Supplementary Table 2**. The columns are drugs standardized with DrugBankID, the rows are P450 enzyme family. Each circle indicates drug targets that belongs to a certain family, presented by HGNC ID. If empty, meaning that drug is absent of targets metabolized that enzyme family. **b.** Forest plot of association of acalabrutinib DDI with drug being metabolized of individual P450 enzymes. **c.** Common adverse event signals associated with predicted DDIs with acalabrutinib in FAERS database**. d.** Distribution of ATC code of predicted DDIs belonging to CYP3 enzyme family. **e.** Distribution of therapeutic subgroup of predicted DDIs.

To characterize common phenotypes underlying aca-DDI drugs predicted to be substrates or inhibitors of CYP3A enzymes, we analyzed individual case safety reports (ICSRs) from FAERS where acalabrutinib was the primary suspect drug. Several common non-fatal side effects were observed across CYP3A-family DDIs when co-administered with acalabrutinib, including headache, fatigue, diarrhea, and malignant neoplasm progression (**Fig. 3c**). We categorized these interacting drugs by their drug classes using first-level and second-level ATC codes. At the main pharmacological group level, nervous system, cardiovascular system, and alimentary and metabolism drugs were frequently reported to have DDIs with acalabrutinib (**Fig. 3d**). At the therapeutic subgroup level, antibacterial and chemotherapy drugs showed frequent DDIs with acalabrutinib (**Fig. 3e**). These findings highlight the diverse range of drug classes that may interact with acalabrutinib, emphasizing the need for proactive DDI prediction across a broad spectrum of co-administered therapies.

### DDI-GPT provides interpretable predictions through gene importance scoring

DDI-GPT generates importance scores for each word in its analysis, revealing which elements most strongly influence its predictions. The evaluation module assesses these scores by introducing small controlled perturbations to input words and measuring how these changes affect DDI predictions (**Fig. 4a, Methods**). Words critical to the prediction show significant changes in DDI probability even with minimal perturbation.

**Fig. 4:**
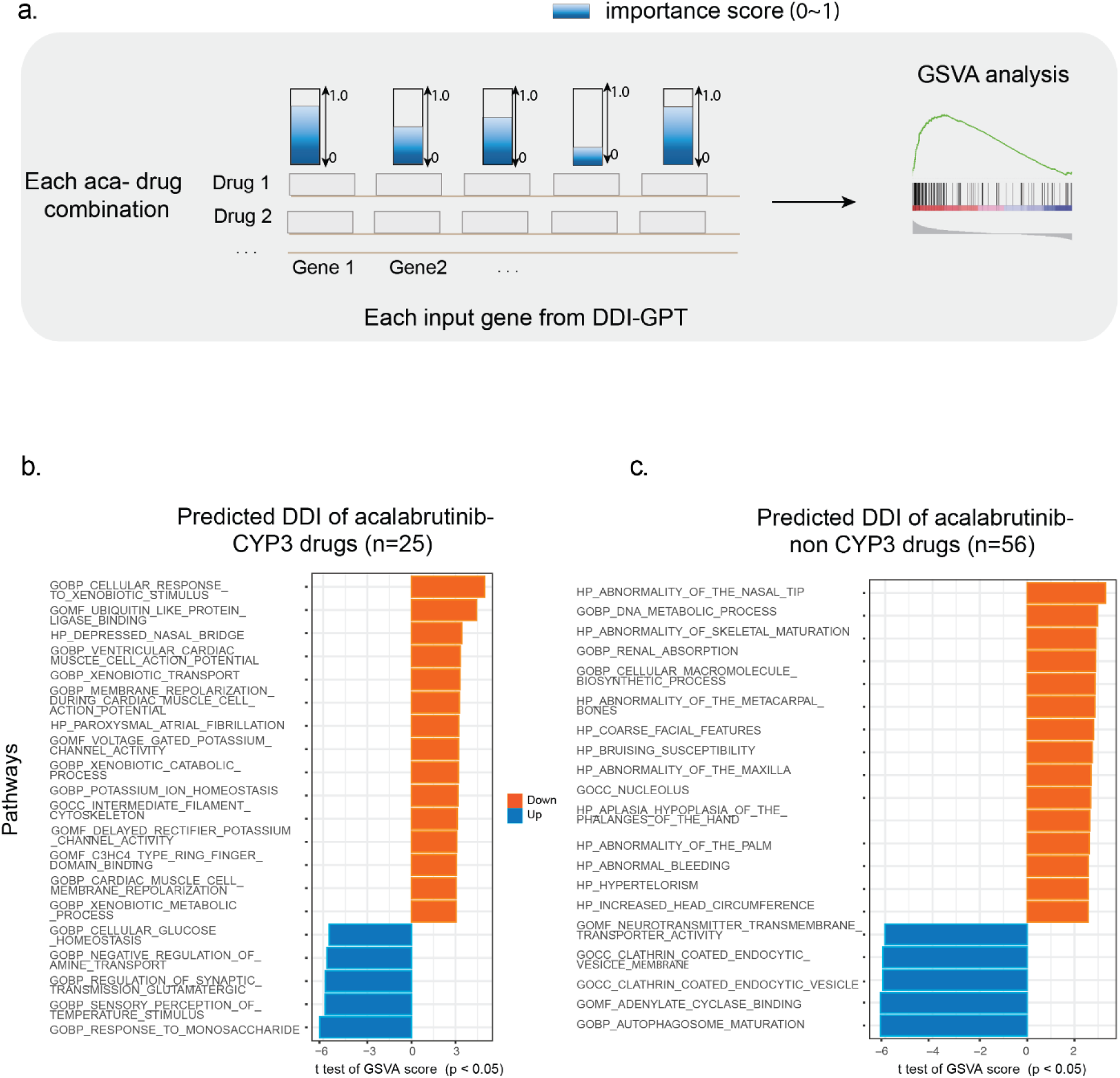
**a.** DDI-GPT evaluation module uses importance scores to estimate the contribution of the word to the prediction, which can be computed efficiently by using perturbation-based approximation. The idea is to perturb the contribution of the word by adding a Gaussian noise and measure the magnitude of change in the prediction score. **b.** Genes are clustered based on their importance value by an unsupervised K-means clustering. Pathway analysis is performed on each cluster.

To identify biologically meaningful patterns, we analyzed importance scores across different acalabrutinib DDIs (aca-DDIs). We hypothesized that genes consistently receiving high importance scores would share biological functions or pathways relevant to drug interactions. After calculating importance scores for each gene across individual aca-DDIs, we performed enrichment analysis using gene ontology (GO) terms, applying Fisher’s exact test with false discovery rate (FDR) correction. The analysis revealed distinct biological pathway associations. CYP3-associated aca-DDIs showed significant enrichment for genes involved in "cellular response to xenobiotic stimulus" and "xenobiotic metabolic process" (q = 2.81 × 10-4, 2.69 × 10-3). This enrichment suggests these interactions primarily involve drugs affecting foreign substance metabolism. In contrast, non-CYP3-associated aca-DDIs showed enrichment for genes in alternative metabolic pathways (**Fig. 4b-c**).

### Network analysis reveals mechanisms underlying predicted DDIs

In our previous study, PAIRWISE predicted 30 drugs as synergistic with acalabrutinib, which we validated through in vitro high-throughput screening^27^. Among these, DDI-GPT predicted 14 drugs (probabilities > 0.5) as having potential DDIs. To understand the mechanisms behind these predictions, we ranked genes by their importance scores and used network analysis to examine relationships between drug target networks and adverse reaction (ADR) networks. We obtained ADR gene sets from the ADReCS-Target database^28^ and constructed drug target networks using DDI-GPT’s highest-ranked genes. For each predicted DDI, we measured the network proximity between genes in the ADR subnetwork and the drug target subnetwork (**Fig. 5a-b**).

**Fig. 5:**
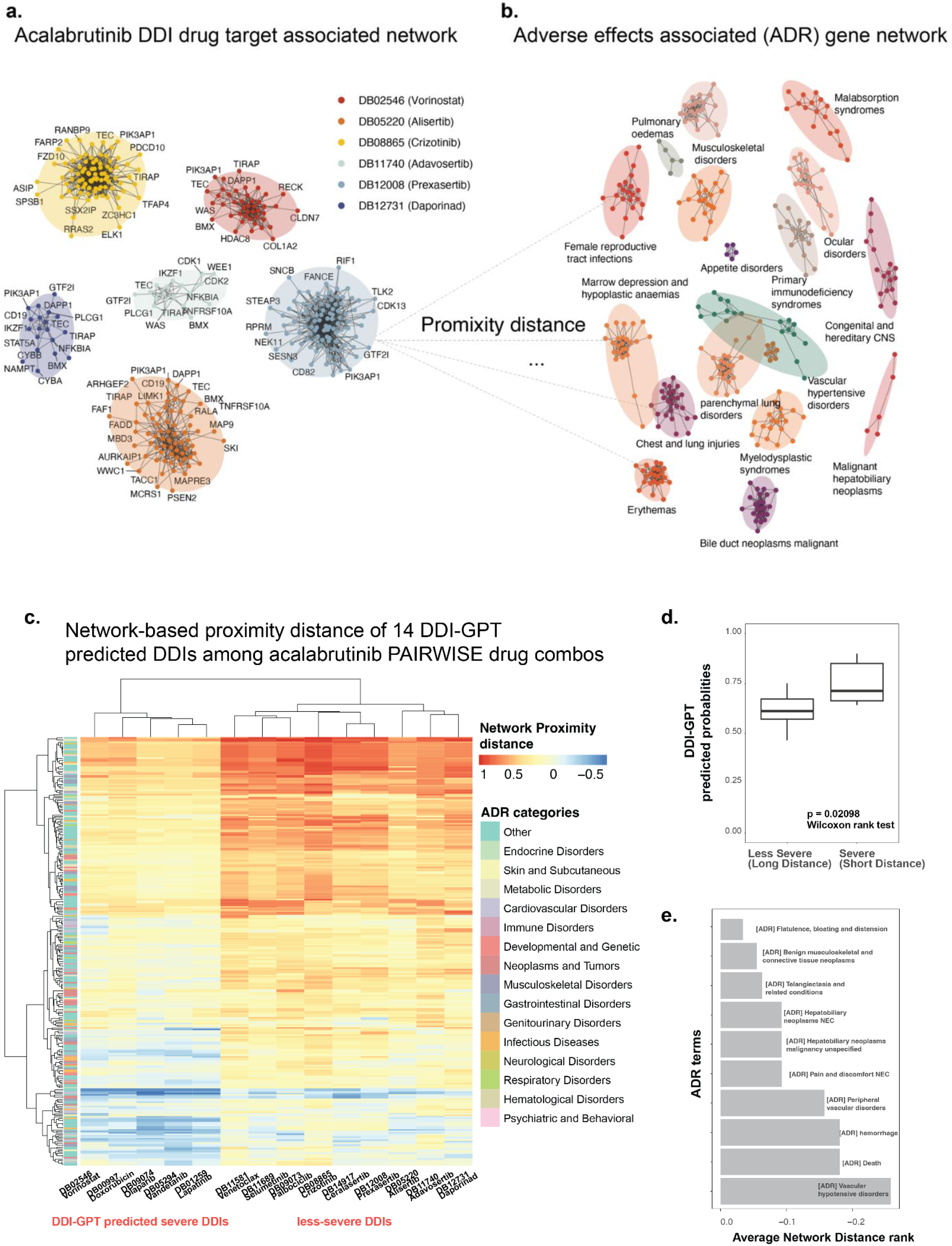
Network distance approach to identify DDI’s gene signatures. **a.** Each cluster in the graph on the left represents a single drug that is predicted as synergistic by PAIRWISE model, and each node within a cluster represents important drug targets ranked by DDI-GPT, with RWR on PPI. **b.** Each cluster in the graph on the right represents genes belonging to an ADR term. **c.** We calculated the network distance between individual drug target subnetwork and each ADR subnetwork. We plotted all combinations with its network distances and unsupervised clustering reveal two distinct DDI profiles among 14 predicted DDIs with the combination involving acalabrutinib. **d.** For 2 DDI profiles (severe or less-severe) created by paring 30 drug target subnetworks and 294 ADR subnetworks, we measured the correlation between the average distances of that profile and the DDI-GPT predicted probability values for drug combinations of that profile. We find that combinations of drugs categorized by distance approach tend to have higher DDI-GPT predicted probability values in severe group (p = 0.02098, Wilcoxon rank test). **e.** The top enriched ADR term in the set of genes contributing to DDI of drug combinations including the acalabrutinib, sorted by their ranking of average network proximity distance across predicted DDIs.

Consistent with our hypothesis that drugs associated with CYP3A family would exhibit aca-DDIs, we found strong CYP3A inhibitors (crizotinib, lapatinib, vorinostat) and moderate CYP3A inhibitors (olaparib, venetoclax, selumetinib) were highlighted by DDI-GPT predictions, as noted by the Center for Drug Evaluation and Research where these drugs require close monitoring when co-administered with acalabrutinib. Remarkably, we identified two distinct sets of drug combinations predicted as DDIs: combinations showing more severe effects when co-administered with acalabrutinib (characterized by shorter network distances) and combinations showing less severe effects (characterized by longer network distances) (**Fig. 5c**).

Measuring the relationship strength between average network proximity distance and DDI-GPT predicted probability scores, we found that the less-severe group marked by shorter average network distance shows significantly increased DDI probability when co-administered with acalabrutinib (p=0.002098, Wilcoxon rank test, **Fig. 5d**). One of the most enriched ADR terms for DDI is "vascular hypotensive disorders" (**Fig. 5e**). The genes associated with this ADR include BTK itself and downstream signaling molecules such as PLCG2 and MAPK1, which play critical roles in vascular homeostasis and endothelial function. Notably, drugs like lapatinib emerge due to their overlapping pathways and targets. For instance, lapatinib is a dual inhibitor of EGFR and HER2 tyrosine kinases and can influence the MAPK cascade—a pathway also modulated by acalabrutinib through BTK signaling. The convergence on the MAPK pathway suggests that co-administration could amplify effects on cell proliferation and vascular responses, potentially increasing hypotensive event risk. Additionally, the enrichment of "hepatobiliary neoplasms" suggests a possible link between acalabrutinib therapy and hepatic cellular processes (**Fig. 5e**). Genes like CYP3A4 and CYP3A5, responsible for acalabrutinib metabolism in the liver, may contribute to hepatotoxicity or neoplastic transformations when their function is altered. Furthermore, the identification of "telangiectasia and related conditions" aligns with vascular anomalies that can arise from BTK inhibition. These findings highlight the utility of network analysis in linking drug target genes to enriched ADR terms.

## Discussions

Here we present DDI-GPT, a framework that seamlessly integrates biomedical knowledge graphs with the semantic capabilities of large language models to predict drug-drug interactions. While previous approaches have achieved high accuracy on benchmark datasets, DDI-GPT demonstrates superior performance in zero-shot predictions on newly curated FDA DDI data. High accuracy alone is insufficient for clinical applications - providing clear explanations to patients, clinicians, and scientists is essential for establishing clinical validity. DDI-GPT combines prediction accuracy with interpretability, offering insights into the complex relationships between DDIs and gene features at both pathway and interactome levels.

Using the TwoSIDES dataset collected before 2013, we developed our DDI prediction model and explored how different knowledge graph configurations and deep learning architectures affect binary DDI prediction. We demonstrate the value of enhancing GPT architecture with a comprehensive knowledge graph that includes both biological process and pharmaceutical property information. Importantly, we explain our complex machine learning model through gene feature attribution methods. DDI-GPT’s evaluation module captures important gene signals highlighted by the model, which are difficult to detect with standard LLMs. These results help identify plausible DDIs for newly approved drugs (such as the associations between CYP3A enzyme family drugs and acalabrutinib). One limitation of our study is that the enzyme family-DDI associations identified by our model cannot definitively establish causation. The presence of CYP3A inhibitors or inducers does not necessarily lead to clinically significant DDI outcomes, as establishing such clinical relevance would require in vivo trials and observations.

We further developed a gene importance score-based pathway enrichment analysis and network distance approach to understand the biological phenotypes associated with DDI-GPT predicted interactions. Our network-distance analysis grouped severe DDIs with adverse drug reaction terms related to vascular homeostasis and hepatobiliary neoplasms, indicating these features share information about DDI severity. Given that FDA-approved drugs from 2013 to 2017 comprised approximately 65% substrates, 30% inhibitors, and 5% inducers of CYP3A^29^, our gene score-based network distance approach helps clinicians select safer drug combinations by identifying high-risk DDIs that show closer proximity to ADR subnetworks.

Our study validation used FDA-curated labels from 2013-2023, representing spontaneously reported DDI records from patients, clinicians and pharmaceutical companies. To evaluate model generalizability, we performed temporal validation on DDI records for drugs not present in the training data. With rapid advances in LLM technologies and automated database annotation projects, a promising future direction is updating knowledge graphs by capturing comprehensive textual information about drug entities from ultra-large databases like PubChem and DrugBank. One limitation is that DDI-GPT cannot reach its full potential for precision medicine when DDI training data lacks personalized information such as transcriptome profiles and gene mutation status. Future performance improvements may come from incorporating *ex vivo* drug toxicity information from PB/PK studies into our framework. Moreover, we envision that real-world data and evidence can further facilitate DDI identification when considering individual tumors in a population^30^.

Knowledge graph-enhanced LLM prediction models will likely increase as graphs are continuously updated and LLM techniques rapidly advance. However, "black box" LLMs without explanations are difficult for clinicians to trust and extract meaningful information from. Therefore, combining complex LLMs with explanation modules for knowledge graphs is both necessary and urgent. DDI-GPT takes an important step toward explainable DDI predictions. This study’s improvements in predictive accuracy and interpretation analysis further strengthen the model’s rationale and clinical utility.

## Methods

Our objective is to predict whether a combination of two drugs has adverse interactions with each other, particularly in a training scenario where limited samples are available. We first explain how we obtain the drug-related information from a heterogeneous knowledge graph (KG). The KG provides LLM with a structured approach to relate concepts. We then describe our proposed method that fine-tunes an LLM-based model for DDI prediction.

### Curation of a Heterogeneous Knowledge Graph

We obtain heterogeneous graph representations of drugs by downloading a large-scale biomedical KG, iBKH (**Supplementary Fig. 1**), which covers all drugs in our benchmark TwoSIDES dataset and the external validation dataset, as well as 4 other types of biology entities, including drug targets, biological pathways, anatomical therapeutic chemicals, and drug categories, as well as the associated relations among them. Our KG is composed of entity-relation-entity triplets. Seven types of relations are used to connect the biomedical entities (**Supplementary table 1**). We also constructed several KGs to represent different prior knowledge of drug entities, to test their effects on model performance (**Supplementary Note 2**).

### Integrating KGs into LLM input

We used a two-step process to utilize an LLM for biomedical graph data: knowledge query and knowledge injection. In the knowledge query step, all the biomedical entities related to a specific drug are selected to retrieve their corresponding triplets. In the knowledge injection step, the queried triplets are injected into sentence at their corresponding position, generating a sentence tree. The structure of the tree is illustrated in **Figure 2a**. In this study a sentence tree can have multiple branches.

#### Knowledge query

To retrieve all relevant information about a drug from its neighbors, we convert the structured graph into text for each instance of a drug. For example, given the biomedical entities (e.g., category, ATC, pathways, target, enzyme, carrier, transporter) and their relations with the input drug “Verapamil”, traditionally we convert the instance as “Drug: Verapamil ATC: C08DA01 Category: Calcium Channel Blocker Pathways: SMP0000375 Targets: CACNA1C Enzyme: CYP3A4 Carrier: ALB Transporter: ABCB1”.

#### Knowledge injection

Direct concatenating a sequence of tokens within a knowledge graph triplet can potentially result in knowledge noise. In general, excessive incorporation of knowledge may divert the sentence from its intended meaning. Originally, K-BERT introduced soft-position and visible matrix to limit the influence (Liu et al., 2019). To overcome these challenges, inspired by K-BERT method, we propose a novel method to integrate KGs into DDI structure data via a sentence tree-based approach. Specifically, the DDI dataset is labeled at the single sentence level. Each sentence has a pair of drugs (*e1, e2*) with an annotated relation. We denote a sentence *s =* {*w_0_, w_1_, w_2_, …, w_n_*} as a sequence of tokens, where n is the length of this sentence. For an input DDI, we first transform it into the sentence “*Drug1* {*RELATION*} with *Drug2*”. The {*RELATION*} is set as a binary label {*interacting, non-interacting*} as ground truth for optimization during the training, and was not seen by DDI-GPT in the input sentence, being replaced with a special MASK token. For an input sentence, the knowledge layer injects relevant triples from a KG into it, transforming the original sentence into a knowledge-rich sentence tree.

### DDI-GPT model architecture

DDI prediction is a task to identify association between drug pairs in an input sentence and assign the right classification to each pair. The task consists of three primary modules: the knowledge module, and the prediction module, and the evaluation module.

#### Knowledge module

The knowledge module takes drug pairs as input, identifies their neighborhood in the KG, and transforms a drug pair into a knowledge-rich sentence tree. This sentence tree is subsequently translated into a token-level embedding representation. Since drug may have long or short dependencies with the other drug and biomedical entities, DDI-GPT adopts self-attention mechanisms to capture the hidden long- and short-dependency information among different entities.

#### Prediction module

The classification module consists of a GPT-2 backbone optimized on a corpus of 500,000 PubMed abstracts and a linear layer for classification. GPT2 is developed with deep structures and pretrained textual data, which is the predecessor of GPT-3 and GPT-4. It achieved state-of-art results on several language modeling tasks, and with the smallest number of parameters (∼350M), which is applicable for fine-tuning with limited computation resources. In this study we applied the GPT-2 pretrained on 500,000 PubMed abstracts, referred to as BioGPT-2. We built DDI-GPT by fine-tuning BioGPT-2 with the TwoSIDES dataset, in order to adjust BioGPT-2 in the context of DDI prediction. We introduced a visibility matrix, which is a binary matrix where biomedical entities belonging to the same drug are visible to each other, however, they are not visible to another drug. i.e., if element *i* is a token of a drug and if element *j* is a token belonging to same drug, M*_ij_* = 1. M*_ij_* =1 if and only if element *i* is visible to element *j*, otherwise M*_ij_* = 0.

To adapt the model for a binary-class classification task, we added a linear layer as a sequence classification on top of the pretrained BioGPT-2, which uses the special [CLS] token at the start of the model output to classify the DDI type. The same tokenizer used in BioGPT-2 was employed in DDI-GPT. Cross-entropy loss was used to optimize the model during the fine-tuning process.

### Ranking important components in DDI-GPT with evaluation module

To quantitatively determine the important input words in DDI-GPT, we adopted a unified information-based measure described by Guan et al^31^. Briefly, a key task in explaining the black-box model is to associate latent representation with the interpretable input units (e.g., words) by measuring the contribution of the inputs. Existing methods largely rely on gradient-based methods; however, these can fail because they cannot explain how information flows through the network. In our study, we explain input importance through perturbation.

We assign a value σi to each input word, where σi is initially a random value between 0 and 1. We then generate noise vectors with the size of input word embedding. These noise vectors are added to the input word embeddings, scaled by the weights specified in σ_i_. This means that σ_i_ indicates how much noise is added to the corresponding input word. Both the original and perturbed input word embeddings are fed into DDI-GPT. We then measure the difference between the two output logits, which indicates how significantly the perturbation affects the outputs.

The value of 1-σ_i_, referred to as 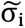, shows how much noise the corresponding input word can tolerate without causing a significant change in the output logits. If a word is important to the outputs, we expect that perturbations to that word’s input embedding will lead to a significant change in the logits. Hence, the reported 1-σ_i_ is proportional to the importance of the words: the bigger the reported 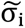, the more important the corresponding input word is.

### Experiments setting

#### Side-effects data

In this study, two real-world datasets were used to evaluate the DDI-GPT performance. For the training, testing and validation of the model, drug side effects are collected from the TWOSIDES dataset. To construct the external validation dataset for assessing the performance of our model, we adopted a new reference set of clinically relevant adverse DDI data^32^. We removed DDIs from this external validation set to ensure there were no overlapping data points seen during the training process. All curated datasets used in our study are provided in the supplementary data.

#### Curation of Newly updated DDI data

We removed drug pairs that overlapped with the training data, yielding 9,480 unique DDIs associated with 442 distinct drugs. To create a negative control set, we randomly paired two drugs from the 442 unique drugs, ensuring the number of non-interacting drug pairs matched the number of interacting DDIs. To further verify that these randomly paired drugs were unlikely to interact, we conducted PubMed queries for each pair and excluded any that returned search results suggesting a known interaction. This rigorous process resulted in a final dataset of 13,586 drug pairs, including both interacting and non-interacting pairs, forming a comprehensive, updated dataset for independent validating DDI predictions.

#### Metrics

Performance was evaluated using area under the receiver-operating characteristic (AUROC), area under the precision-recall curve (AUPRC) and average precision at 50 (AP@50). Higher values indicate better performance.

#### Evaluation strategies

We used the same data splitting procedure introduced by Zitnik et al^14^. For every experiment, we conduct five independent runs with different random splits of dataset. We then selected the best performing model based on AUROC performance from the validation set. The selected model was then evaluated on the test set.

### Disproportionality analysis to determine CYP enzyme families associated with enriched DDIs predicted by DDI-GPT

We propose using Enzyme Odds Ratio (EOR) to evaluate the possible association between a specific Cytochrome P450 (CYP) enzyme family and the DDIs predicted by DDI-GPT This approach allows us to investigate whether a specific CYP enzyme family is enriched in DDIs more than would be expected by chance. There are 17 CYP enzyme families.

The EOR compares the odds of a specific CYP enzyme being associated with a particular DDI drug pair to all other enzymes, relative to the reporting odds for other drugs. We performed a disproportionality analysis to calculate the EOR. Before conducting this analysis, we first created a CYP enzyme contingency table (**Supplementary Table S2**) which served as the basis for the subsequent calculation of the EOR. The following formulas were used to calculate the EOR and its 95% confidence interval (CI):

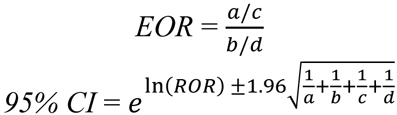

In these formulas, *a*, *b*, *c*, and *d* represent the counts from the contingency table. For the disproportionality analysis, we used the Chi-square test or Fisher’s exact test, and all statistical tests were two-tailed. All statistical analyses were conducted using Python SciPy v1.11.2 software.

### DDI-specific pathway analysis

To analyze the biological processes relevant to DDIs containing specific drugs in the dataset, we tested the top-ranked genes in the lists ordered by the importance values generated by the perturbation approach. We calculated pathway enrichments using GeneSet Variation Analysis (GSVA) R software package^33^ based on the importance value matrix for all predicted DDIs and non-DDIs containing the drug acalabrutinib. We used the set of pathways from Gene Ontology (GO) Biological Process terms for enrichment tests. The enrichments were calculated by a non-parametric estimator implemented in the ‘gsva()’ function. False discovery rate (FDR) correction was applied using the Benjamini-Hochberg procedure.

### Network distance measures

The adverse drug reaction (ADR) gene set was retrieved from ADReCS-Target (Huang et al., 2018), totaling 294 ADR phenotypes across 55,340 pairwise gene-ADR associations. The drug target gene set was retrieved from DrugBank database v 5.1.0^34^. A PPI network was obtained from STRING database v12.0^35^. We performed a Random Walk with Restart (RWR) algorithm to explore the global network structure of drug target on PPI interactome, using drug target as seed nodes to visit all nodes in PPI. The visiting nodes with higher-probabilities were used to complement and construct the drug target subnetwork. We employed network-based distance measures proposed by Cheng et al., (2019). In our context, given A, the set of ADR genes, T, the set of drug targets, and *d(A,T),* the closest distance measured by the average shortest path length between each node *α* ∈ A and its nearest drug target *t* ∈ T in PPI, the distance is defined as:

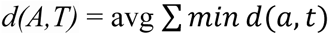

Here, *d(a,t)*represents the shortest path length between nodes *α* and *t*.

### Development of standalone application

To support practical real-world application and assist domain experts in exploring DDIs predicted by DDI-GPT, we have developed an interactive tool. This tool is a Python application coupled with the neo4j database that leverages our KG features at the backend. The frontend (i.e., the web application portal) was built using Streamlit v1.38.0, and NetworkX v.3.2.1 was used for graph visualization and data exploration. It is designed to support clinical decision-making and patient safety management. Users can search by entering a combination of two drugs, after which the backend framework automatically transforms these inputs into biologically meaningful sentences for DDI-GPT prediction. Our pre-trained DDI-GPT model then generates predictions for each DDI. The application also displays protein-protein and side-effect networks, which are visually represented as relational networks. Additionally, the application provides a table containing k-shortest path inference information from one to the other drug, as illustrated in the complex network, to facilitate analysis.

## Code availability

DDI-GPT and the benchmarking platform can be downloaded from GitHub https://github.com/Mew233/ddigpt. Visualizations of DDI-GPT web server can be accessed on https://pyvisddi-24u28afk4upfhpvclyujvs.streamlit.app/

## Acknowledgement

The project was funded by AstraZeneca. O.E. is supported by UL1TR002384, UG3CA244697, R01CA194547, P01CA214274 grants from the National Institutes of Health and LLS SCOR grants 180078-02, 7021-20, 180078-01. H.P. was sponsored by Beijing Nova Program (20220484073). The authors would also like to thank the wider collaborative teams from Weill Cornell Medicine and AstraZeneca for their support and valuable feedback right through the project.

## Supplementary materials

**Supplementary Table 1.** Our curated KG is heterogenous, with 4 types of entities and 7 types of undirected edges. Below tables shows a breakdown of entities by entity type and relations by entity type. Supplementary Note 1 provides detailed information about definition of entities.

| Entity pair | Relation types | Number of relations |
| --- | --- | --- |
| Drug-Gene | Drug-Target-Gene | 16,518 |
|  | Drug-Transporter-Gene | 3,066 |
|  | Drug-Enzyme-Gene | 5,241 |
|  | Drug-Carrier-Gene | 853 |
| Drug-Pathway | Drug-Association-Pathway | 3,231 |
| Drug-Category | Drug-Belongs to-Category | 599 |
| Drug-ATC | Drug-is_a-ATC | 599 |

**Supplementary Table 2.**
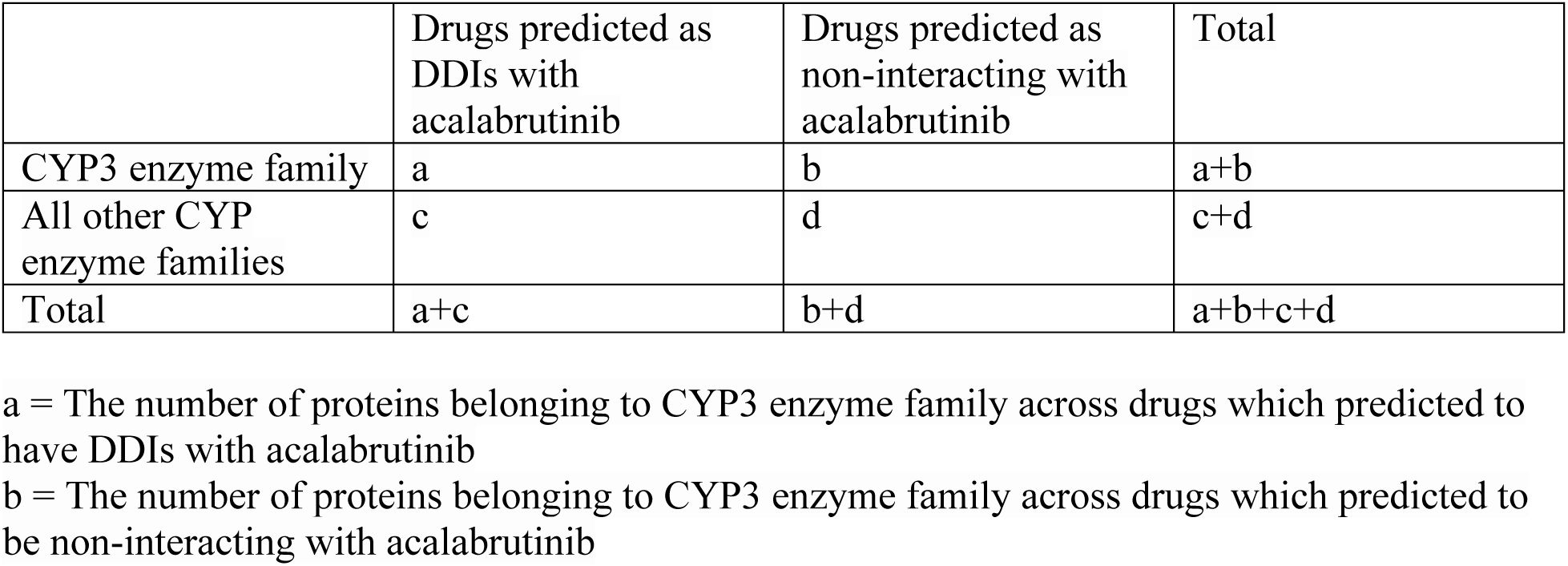
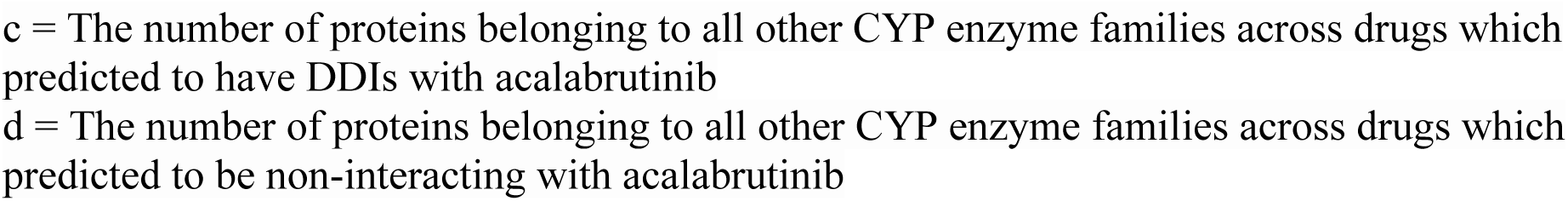
Contingency table used for calculating EOR.

**Supplementary Table 2.** KG features are critical for model performance. On limited KG, DDI-GPT still achieves robust performance compared to CNN and RNN.

| Method | Drug features injected from biomedical Graph | Architecture | AUROC | AUPRC | Precision@50 |
| --- | --- | --- | --- | --- | --- |
| DDI-GPT<br>(ablated KG with biological process alone) | pathway, targets | GPT framework | **0.910±0.005** | **0.835±0.003** | **0.829±0.003** |
| CNN<br>(ablated KG with biological process alone) | pathway, targets | Convolutional Networks | 0.762±0.002 | 0.601±0.005 | 0.702±0.004 |
| RNN<br>(ablated KG with biological process alone) | pathway, targets | Recurrent Neural Networks | 0.574±0.001 | 0.395±0.004 | 0.660±0.003 |
| DDI-GPT<br>(ablated KG with Pharmacological properties alone) | ATC, category | GPT framework | **0.938±0.002** | **0.873±0.003** | **0.852±0.006** |
| CNN<br>(ablated KG with Pharmacological properties alone) | ATC, category | Convolutional Networks | 0.815±0.003 | 0.683±0.004 | 0.734±0.002 |
| RNN<br>(ablated KG with Pharmacological properties alone) | ATC, category | Recurrent Neural Networks | 0.577±0.004 | 0.401±0.002 | 0.670±0.003 |

### Supplementary Note 1

- *category*: This relation links drug nodes to MeSH (Medical Subject Headings) nodes.
- *ATC*: This relation links drug nodes to ATC (Anatomical Therapeutic Chemical) classification system code nodes.
- *pathway*: This relation links drug or protein nodes to pathway nodes.
- *target*: This relation links drug nodes to protein nodes.
- *enzyme*: This relation links drug nodes to protein nodes that catalyze chemical reactions involving the drug.
- carrier: This relation links drug nodes to protein nodes that are secreted proteins binding to drugs and carrying them to cell transporters.
- *transporter*: This relation links drug nodes to protein nodes representing membrane-bound proteins that shuttle ions into or out of cells.

### Supplementary Note 2: Choice of graph to represent prior knowledge of the drug entity

We decided to proceed with the iBKH network to represent prior knowledge about the drug entities of interest when predicting co-prescribed drug combinations. The choice was made because iBKH network had the best coverage of the drug molecules we were interested in, and was the most general-purpose for application to future tasks. We studied the impact of varying the components of the curated KG that are used as prior knowledge.

#### Varying KG components

We first studied the impact of varying the components of the KG used as background knowledge for knowledge injection into the DDI sentence. We found that removing biological process-related entities, such as drug target and pathways, resulted in most significant decrease in performance This fits our assumption that drug targets are often critical for DDI prediction. Genes that share similar biological pathways should result in similar predicted DDI risk. While molecular function often overlaps with biological process annotation, biological process terms are more diverse and represent specific biological end states or outcomes. There was minimal impact on performance when pharmacological properties, such as ATC codes and categories, were removed.

#### Varying resolution of KG

Given a source drug, we query the KG by identifying its drug targets, transporters, carrier proteins, with the maximum of *k = 2* of each node type. These *k* proteins are then connected to source drug in the KG. By varying the *k* parameter, we can study the influence of the density of edges in the graph. We observed the value of *k =2* resulted in best performance of DDI-GPT. When we reduced the value to *k =1*, thereby decreasing the density of the graph, we observed that the performance of DDI-GPT decreased. Increasing the value to *k =3* or more had minimal impact on performance. We assume that genes that share similar biological pathways should result in similar predicted DDI risk.

**Supplementary Fig. 1:**
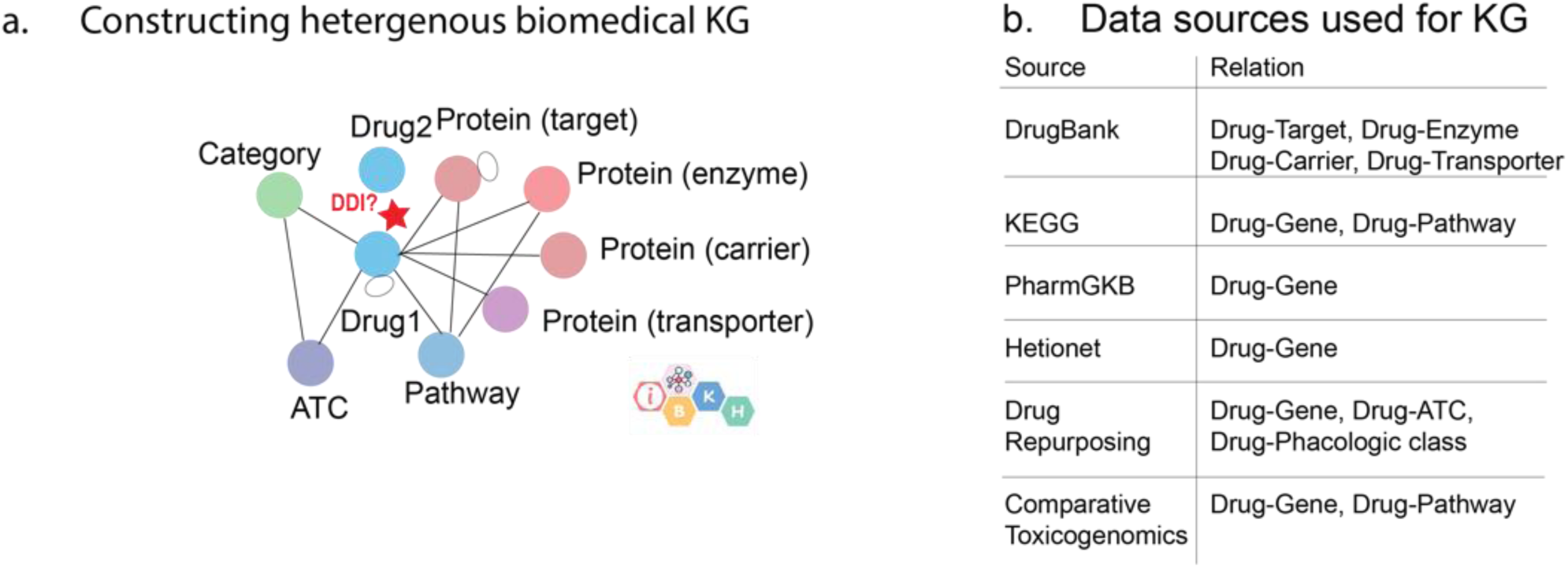
**a.** A schema of drug-related KG curated from iBKH. Each circle denotes an entity type, and each link denotes a relation between a pair of entities. **b.** Data sources integrated for constructing drug-centered KG.

**Supplementary Fig. 2:**
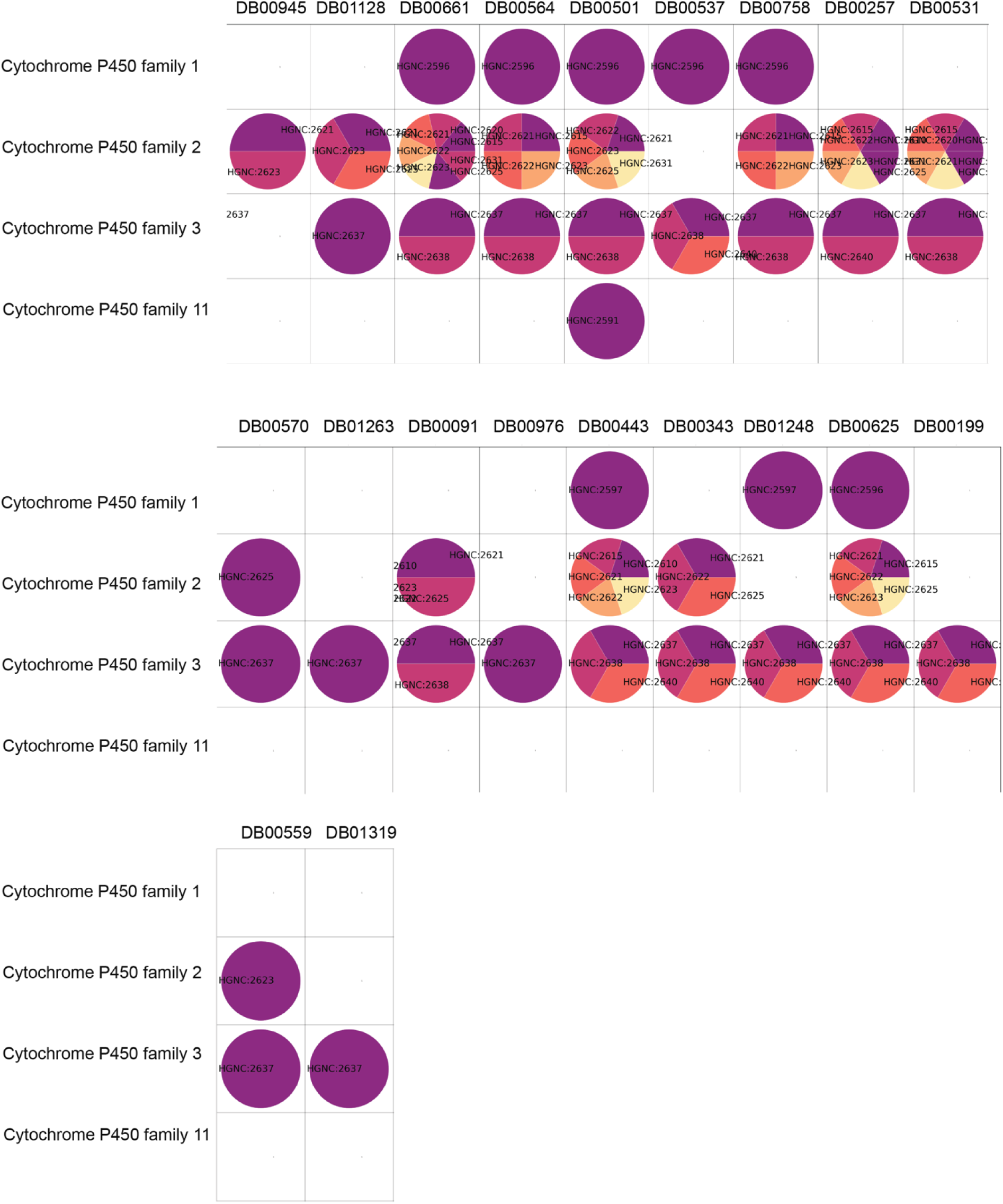
Enzymes family of acalabrutinib predicted DDI drug for disproportional analysis in **Supplementary Table 2**. The columns are drugs standardized with DrugBankID, the rows are CYP enzyme family. Each circle indicates drug targets that belong to a certain family, presented by HGNC,ID. If empty, meaning that drug are absent of targets metabolized that enzyme family.

**Supplementary Fig. 3:**
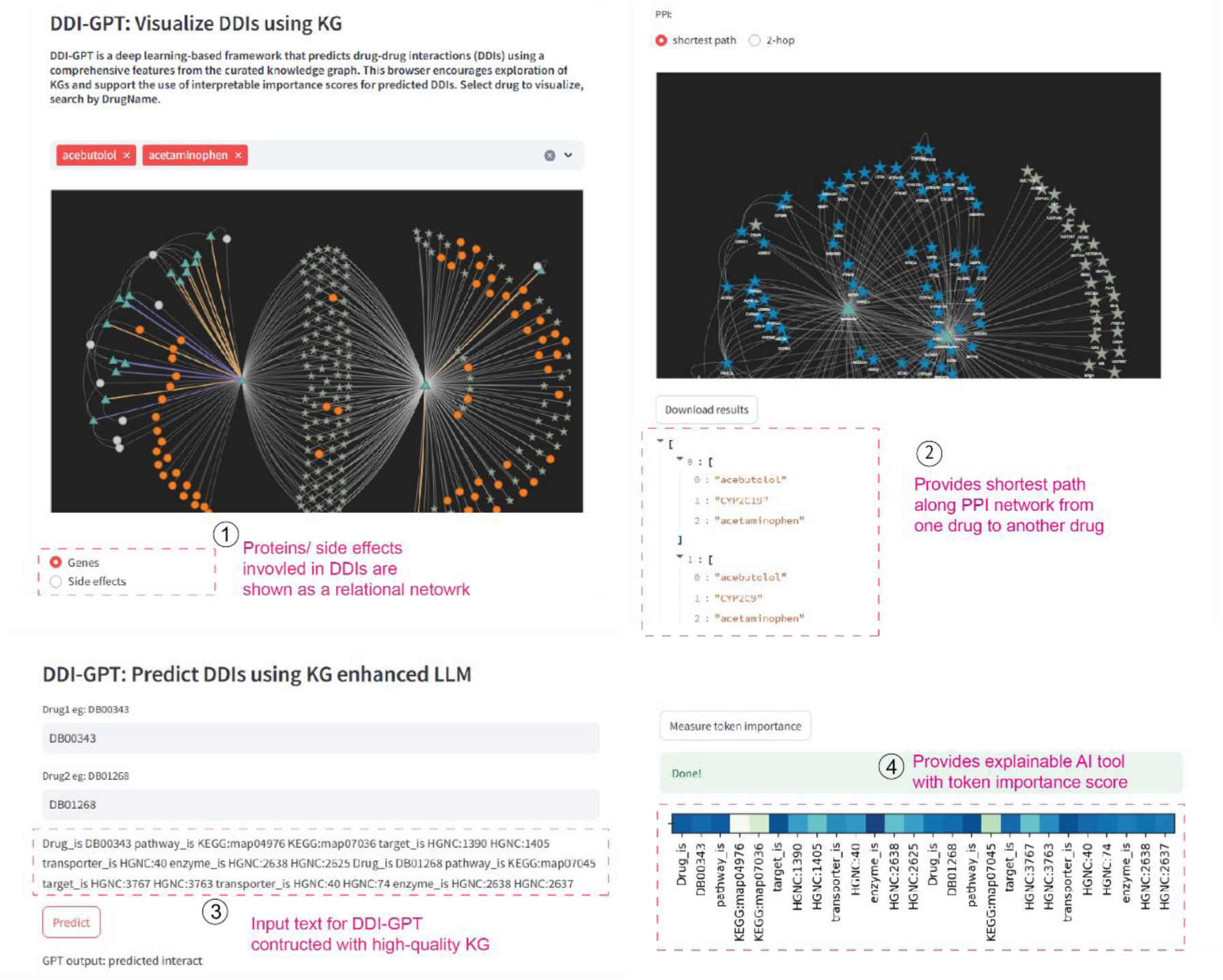
DDI-GPT web server for knowledge-enhanced DDI exploration**. 1.** The user submits a pair of drugs, referenced by DrugBank ID, the input sentence is generated by incorporating drug-related biomedical entities from KG. The input is then set to a cluster running the prediction pipeline, where it is processed. **2.** Once the interface prediction is complete, the DDI-GPT web server will be updated to visualize importance words generated by explanation module. **3.** DDI-GPT web server allows users to explore the KGs that were used by DDI-GPT framework. as well as various drug-protein-drug interaction networks and drug-side-effects-drug interaction network. **4.** The user can view shortest paths or two-hop paths between the combination of drugs.

